# Transcriptomic analysis of TGFβ-mediated fibrosis in primary human Tenon’s fibroblasts

**DOI:** 10.1101/2024.03.09.583791

**Authors:** Zoe Pasvanis, Antony Boynes, Roy C.K. Kong, Elsa C. Chan, Raymond C.B. Wong, Jennifer Fan Gaskin

## Abstract

Glaucoma filtration surgery (GFS) is performed to slow down disease progression in glaucoma, a leading cause of irreversible blindness worldwide. Following surgery, pathological wound healing may lead to conjunctival fibrosis and filtering failure. Myofibroblasts are the key cells responsible for postoperative conjunctival scarring. This study aims to further understand the molecular mechanisms of conjunctival fibrosis following GFS. We utilised RNA-sequencing (RNA-seq) to delineate the TGFβ1 induced changes in the transcriptome of human Tenon’s fibroblasts (HTFs). RNA sequencing was performed on HTFs after 5 days of TGFβ1 treatment. Following quality control, 3,362 differentially expressed genes were identified, of which 1,532 were upregulated and 1,820 were downregulated. We identified signaling pathways associated with the pathogenesis of conjunctival fibrosis. The DEGs (differentially expressed genes) were enriched in pathways including myofibroblast differentiation, TGFβ-signaling, collagen and extracellular matrix organization, epithelial to mesenchymal transition, and cell cycle regulation. The results of this study identified the transition from HTF to myofibroblast is characterised by the upregulation of key genes including *LDLRAD4, CDKN2B, FZD8, MYOZ1*, and the downregulation of *SOD3, LTBP4* and *RCAN2*. This unprecedented insight into the transcriptional landscape of HTFs and myofibroblast differentiation is essential to understand the pathophysiology of conjunctival scarring and develop new therapeutic agents.

## Introduction

Glaucoma filtration surgery (GFS) is the gold-standard surgical procedure performed to lower intraocular pressure (IOP) in glaucomatous eyes. GFS allows for a gradual egression of aqueous humor from the anterior chamber of the eye into a surgically created subconjunctival space, resulting in the formation of a filtering bleb. As a result of surgical intervention, the body’s natural wound healing response is activated. In some cases, the magnitude of this response can be excessive, such that pathological wound healing is observed. This can lead to the postoperative complication of conjunctival fibrosis and surgical failure. Despite the use of adjunctive antimetabolite agents to control scarring, the operation still has a 50% failure rate over 5 years ^1^. Moreover, the standard-of-care antimetabolite agents are nonspecific, cytotoxic cancer drugs such as Mitomycin C (MMC) that can result in toxicity of surrounding tissue, long-term wound breakdown and predispose the eye to severe infections that lead to blindness and even loss of the eye ^2^ . Therefore, there is an unmet medical need to develop novel anti-scarring strategies.

One of the key effector cells responsible for postoperative conjunctival scarring is the human Tenon’s fibroblast (HTF). In an environment of injury and inflammation, as can be produced by GFS, HTFs are activated via TGFβ signalling. As a result, HTFs change functionally and phenotypically into myofibroblasts, which are specialised contractile fibroblasts that produce excess ECM components such as collagens and fibronectin ^3^. Myofibroblasts are typically characterised by microfilament bundles expressing high levels of alpha smooth muscle actin (αSMA) that are organised into stress fibres; they are also vital for mediating contractile function as part of the wound healing response^4^. While myofibroblasts are a vital component of physiological wound healing, an abundant accumulation at the surgical site is one of the major causes of surgical failure^2^. Interestingly Jeon et al. have recently shown that remodelling myofibroblasts to a fibroblast-like phenotype reduces fibrosis in a feline model of established corneal scarring ^5^, hence targeting myofibroblasts by manipulating phenotypic changes may well be an effective way to limit fibrosis in post-GFS scarring. Since there has been no study investigating the molecular mechanism of fibroblast-to-myofibroblast transition by HTF, the aim of the study is to establish the genomic profiles of TGFβ1-induced conversion of HTF to myofibroblasts using RNA-sequencing (RNA-seq).

## Results

### Transition of HTF to Myofibroblast

Using primary HTF, we first performed time-point analysis to determine the peak expression of the marker of myofibroblasts, αSMA. We treated HTFs, derived from 3 GFS patients, with TGFβ1 for 3-6 days (Figure 1). Following real time qPCR, expression of *ACTA2* (which encodes αSMA) in TGFβ groups was significantly increased when compared to control at Day 4 & Day 5 (Figure 2a). In addition, we observed morphological change of HTFs into myofibroblasts (Figure 2b, Supplementary figure 2), where presence of αSMA (red) in collagen fibres is more pronounced, most noticeably occurs at Day 5. Collectively this data suggests that peak activity of HTF to myofibroblast transition occurs five days after treatment with TGFβ.

**Figure 1:**
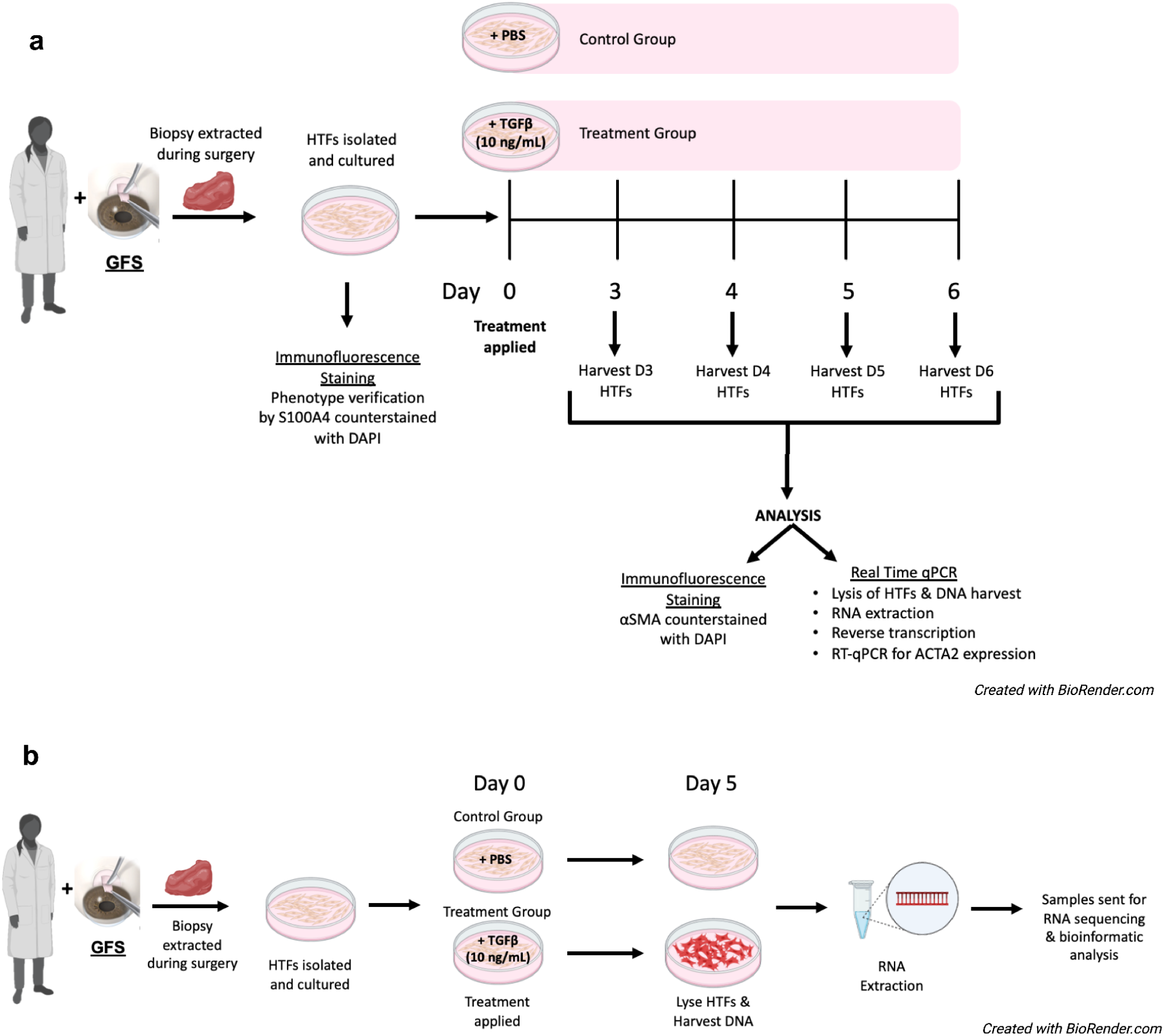
Overview of Project Design. **(a)** Preparation and treatment schedule of in vitro primary human Tenon’s fibroblasts for myofibroblast activation time-point determination (n=3). **(b)** Preparation and treatment schedule of in vitro primary human Tenon’s fibroblasts for RNA sequencing (n=3).

**Figure 2:**
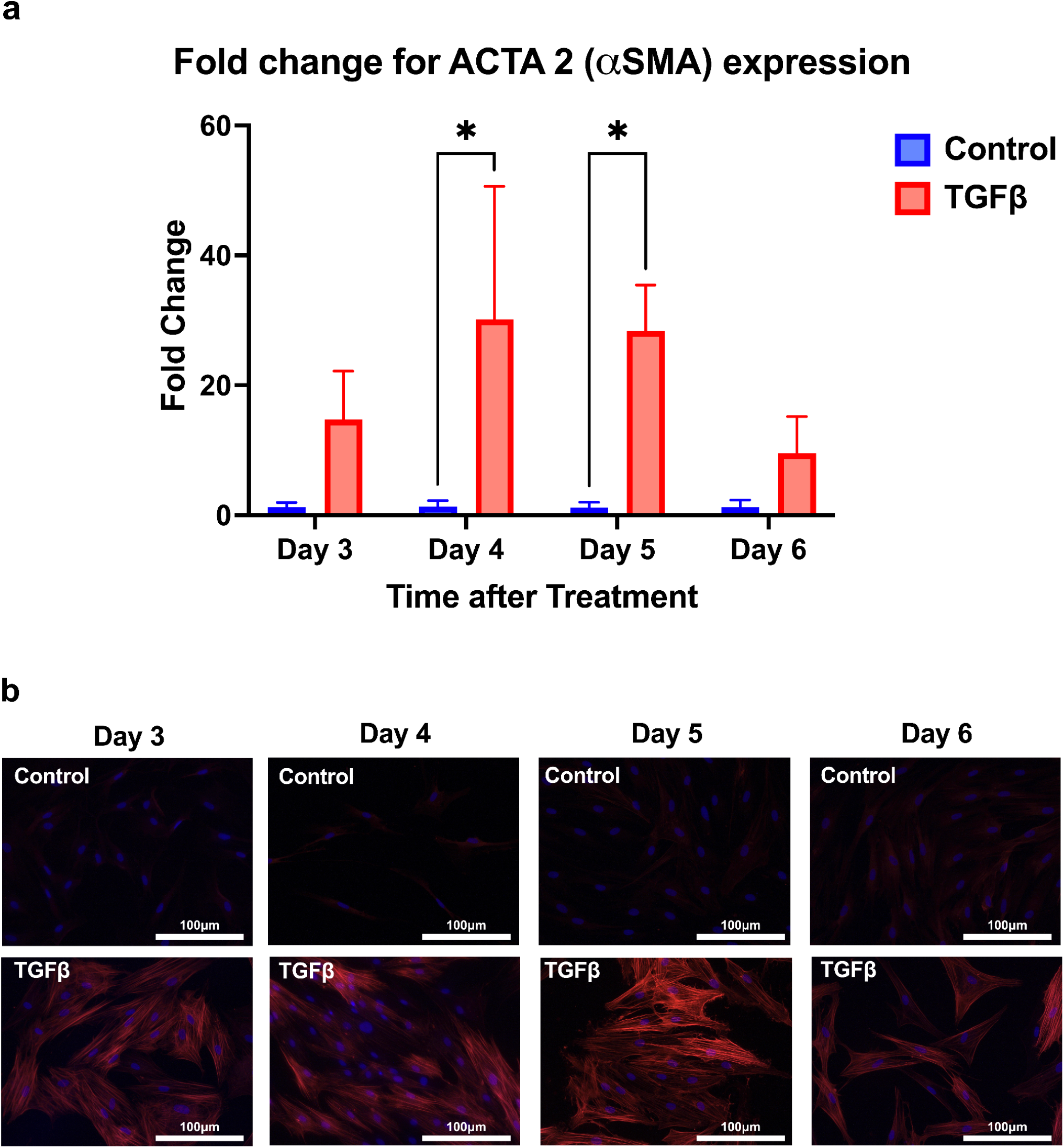
Assessment of myofibroblast activation time-point following TGFβ stimulation. **(a)** Fold change in expression of gene ACTA2 (encoding αSMA) in HTFs over 3-6 days following control and TGFβ treatment, n=3. *\* indicates significance (P<0.05, one-way ANOVA).* **(b)** HTF cell morphology 3-6 days following control and TGFβ treatment. αSMA stress fibres of myofibroblasts (red) identified by immunofluorescence microscopy (magnification 20x) with use of anti-αSMA and fluorophore-labelled secondary antibody. Nuclei counterstained with DAPI (blue).

### RNA-Seq analysis of myofibroblast transition

Since the peak phenotypic changes of fibroblasts to myofibroblasts occurred at day 5 following treatment with TGFβ1, we performed RNA-seq on HTFs from both control (CX2202, CX2203, CX2207) and 5-day TGFβ1 (TGF2202, TGF2203, TGF2207) treatment groups. We first explored the similarity of our samples using principal component analysis (PCA) and hierarchical clustering. Both analyses demonstrate a significant treatment effect of TGFβ1 (Figure 3). PCA analysis presents the variability within the expression data set according to principal components, where we observed biological repeats of the TGFtreated group or control group cluster together, supporting similarities between biological repeats (Figure 3a). Similarly, hierarchical clustering map displays a correlation of gene expression for all pairwise combinations of biological repeats of the treatment group. Samples having correlation values of >0.9 suggest there is no outlying sample (Figure 3b). Together these plots suggest that the data are of good quality, and that it is appropriate to proceed with further differential expression analysis.

**Figure 3:**
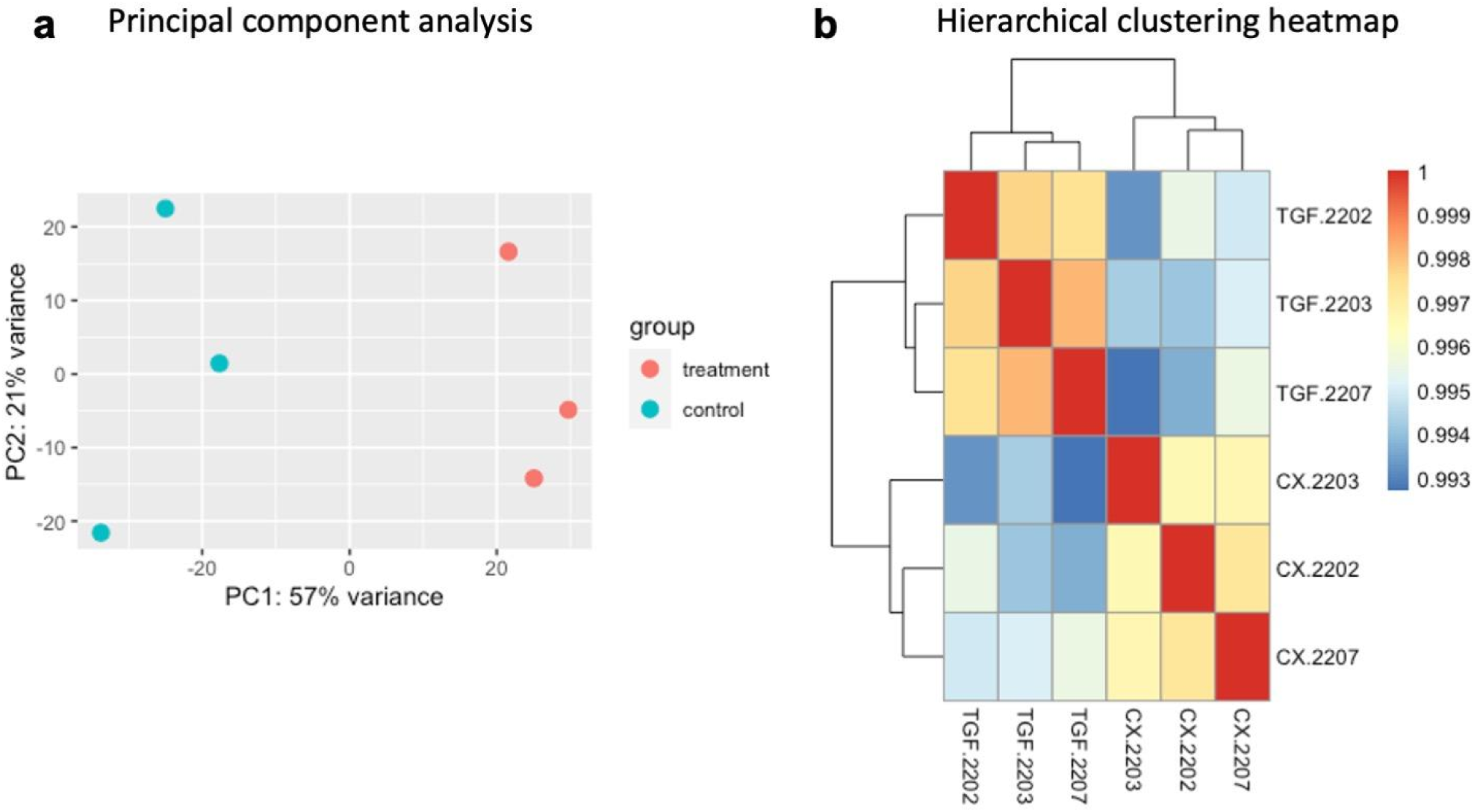
Assessment of TGFβ treatment group variability. **a)** Principal Component Analysis (PCA) plot displaying 6 samples along PC1 and PC2, which describe 57% and 21% of the variability within the expression data set respectively. PCA was applied to data following rlog transformation of normalised counts using DESeq2. Treatment group specified by symbol colour, blue = untreated control HTFs, red = TGFβ1-treated HTFs. **(b)** Hierarchical clustering map displaying correlation of gene expression for all pairwise combinations of samples. Hierarchical clustering was applied to data following rlog transformation of normalised counts (units) using DESeq2. CX=control and TGF=TGFβ1 (10 ng/mL for 5 days).

### Differential Gene Expression

To investigate the impact of TGFβ on HTFs activation into their myofibroblastic phenotype, we performed RNA-seq on primary HTFs. A total of 35,691 genes were identified in all samples. Following DESeq2 analysis and hierarchical clustering, there were 3,362 differentially expressed genes (DEGs) identified, of which 1,532 were upregulated and 1,820 were downregulated in HTF following TGFβ1 treatment (with an adjusted p value of <0.05, Figure 4a). The top 20 DEGs between the two groups are visualized in both a dotplot (Figure 4b) and volcano plot (Figure 4c). The normalised dot plot visualises the differential expression of the top 20 genes in our gene set by relative position (determined by log10 normalised counts) of TGFβ treatment group sample dots (blue, pink and teal) in comparison to control group sample dots (red, yellow and green) within each column. Blue, pink and teal dots located above red, yellow and green dots indicates upregulated expression of the gene in the TGFβ treatment group, whereas the inverse indicates downregulated expression of the gene in the TGFβ treatment group (Figure 4c). Our results indicated that TGFβ1 treatment in HTF upregulated a number of genes including *LDLRAD4, SCUBE3, TXNDC5, CDKN2B, DACT1, DYNC1I1, FZD8, FOXP4, PXDC1, MYOZ1, LANCL2, FBLN5,* while downregulated genes include *IGSF10, SVIL, SECTM1, LTBP4, TNXB, RCAN2, CXCL12* and *SOD3*.

**Figure 4:**
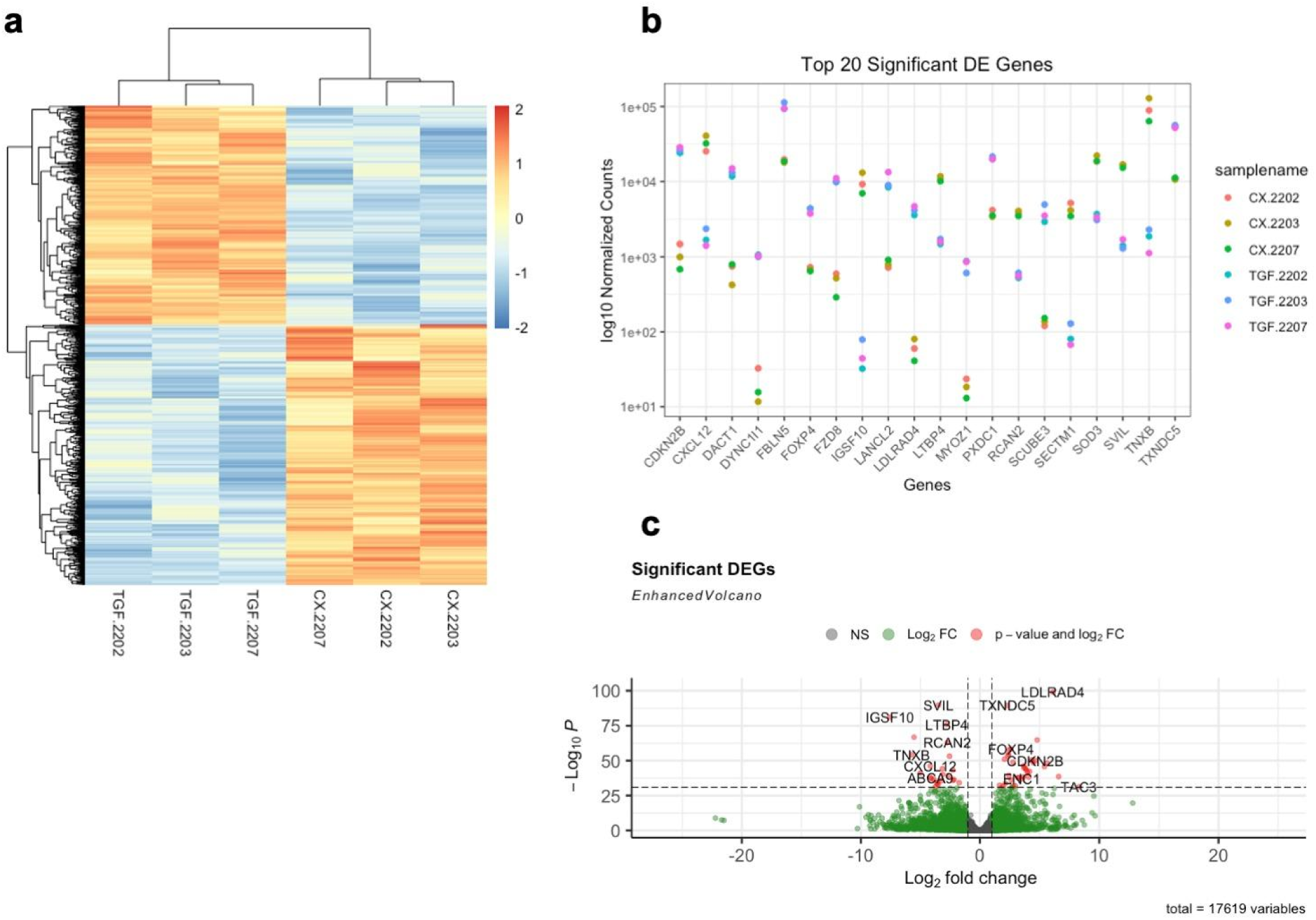
Identification of differentially expressed genes (DEG) in HTF following TGFβ1 treatment. **(a)** Hierarchical clustering DEG heatmap by sample and transcripts on all significant genes using normalized counts **(b)** Dot plot of top 20 DEGs by p adj values of 0.05 inclusive of up and down regulated genes ordered alphabetically. **(c**) Volcano plot with a default log_2_ fold change value of >2 or <-2 and 10e^-32^ p value cut off. Red dots on the right side of the plot indicate genes with significantly upregulated expression, while red dots on the left side of the plot indicate genes with significantly downregulated expression.

### TGFβ Promotes Cell Cycle Arrest in HTFs

TGFβ is known to cause cell senescence in many cell types but may up or downregulate cell proliferation, and cell death in others ^6^. We investigated cell cycle regulatory genes to further understand their role in TGFβ-mediated HTF differentiation or activation. We identified several cell cycle related gene sets from the Molecular Signatures Database as well as our own laboratory dataset ^7^. Tumour suppressor genes such as *CDKN2B* were highly upregulated (Figures 4b and 4c) while *TMPO*, a cell cycle proliferator, was upregulated in the TGFβ treatment group (Figure 5).

**Figure 5:**
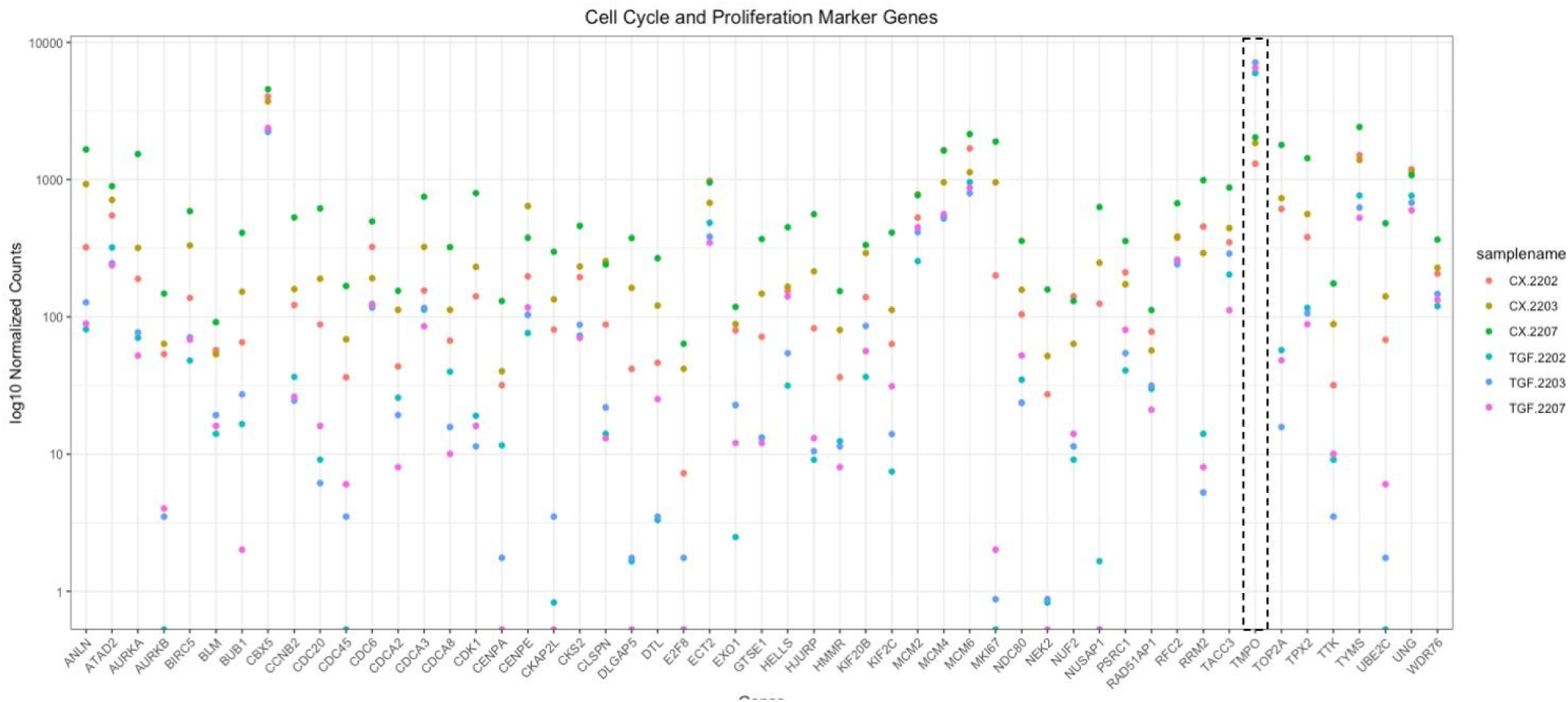
Cell cycle and proliferation gene expression in HTF following TGFβ1 treatment. Cell cycle and proliferation marker genes. Dot plot of normalized counts for known cell cycle and marker gene expression in control (CX2202, CX2203, CX2207) and treatment groups (TGF2202, TGF2203, TGF2207) by biological replicate with padj cutoff of 0.01. All genes were downregulated in the treatment group aside from TMPO which was upregulated (identified by dashed rectangular outline).

### Gene ontology and functional analyses

Next, we performed gene ontology (GO) analysis to understand the functions of the DEG following TGFB1 treatment in HTFs. GO enrichment analysis results revealed top significant enrichment pathways were processes involved in fibrosis including extracellular matrix (ECM) organization, external encapsulating structure organization, collagen fibril organization, and actin filament bundle assembly (Figure 6a, 6b, and 6c). Interestingly our results showed TGFβ regulates several signaling pathways such as Smad phosphorylation, Wnt signaling pathway, and response to cytokines (Figure 6b and 6c). These findings suggest multiple profibrotic pathways in the fibroblast to myofibroblast transition and confirm previous findings that Wnt pathway and Smad signaling promotes fibroblast to myofibroblast transition potentially through cytokines such as IL-11^8^. Regulation of the cell cycle process and regulation of the immune response were highly enriched terms suggesting TGFβ signaling controls cell proliferation and survival (Figure 6a). Cellular component terms reveal interesting insight into the molecular processes mediated by TGFβ. The majority of molecular functions suppressed by TGFβ were found to occur within the cell nucleus and be involved in processes including regulation of cell cycle process and mitotic cell cycle process. Conversely, the activation effects on peptidyl-proline 4-dioxygenase activity (Figure 6a and b) and regulation of collagen occur within the collagen network (Figure 6d and 6e). These findings suggest that fibroblast to myofibroblast transition is regulated by the ECM microenvironment and TGFβ promotes cell senescence through transcriptional induction of the expression of cell-cycle inhibitors.

**Figure 6:**
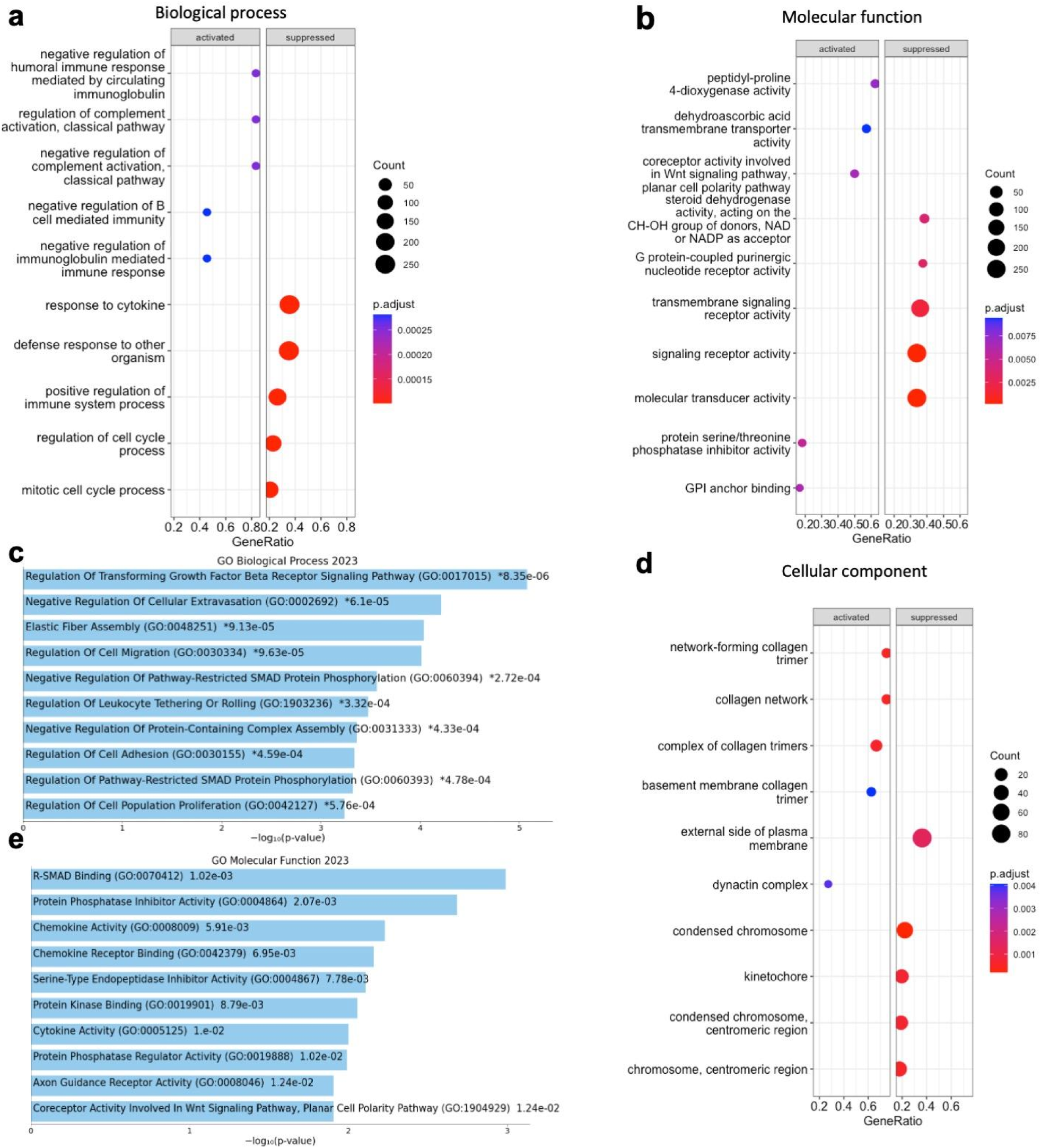
Gene ontology analysis of HTF following TGFβ1 treatment. GO and KEGG pathway enrichment analysis TGFβ1 treatment group compared to control. **(a)** Dotplot shows the activated (upregulated) and suppressed (downregulated) GO terms of biological function associated with all DEGs. The size of the dot is based on gene count enriched in the pathway, and the colour of the dot shows the pathway enrichment significance. **(b)** Dotplot shows the activated (upregulated) and suppressed (downregulated) GO terms of molecular function associated with all DEGs. **(c)** Bar chart of top 10 enriched GO biological function for top 50 DEGs ordered by p-value. **(d)** Dotplot shows the activated (upregulated) and suppressed (downregulated) GO terms of cellular component associated with all DEGs. (e) Bar chart of top 10 enriched GO molecular function for the top 50 DEGs ordered by p-value.

KEGG enrichment analyses revealed top enrichment pathways included TGFβ signaling pathway, Wnt signaling pathway, cell adhesion molecules and NF-kappa B signaling pathway. BMP, LTBP1, Smad Anchor for Receptor Activation (SARA) and SMAD4 were shown to play important roles in the TGFβ-signaling pathway (Figure 7a). Genes from the Frizzled and Smad families were found to be involved in the Wnt-signaling pathway. SMAD3 and SMAD4 specifically were found to play a role in TGFβ and Wnt-pathway mediated cell cycle arrest (Figure 7b). Cell adhesion molecules showed minor downregulation for the majority of cell types such as macrophages, T cells, and B cells. However, there was significant upregulation of myoblasts cell adhesion molecules predominantly driven by Cadherin-2 (CDH2) (Figure 7c). Cadherin expression and cell to cell adhesion play a critical role in transition between cellular states. Here we show CDH2 as a key mediator of intercellular adhesion between myofibroblasts. KEGG network topology showed the crossover between TGFβ-signaling and Smad pathway signaling by LEFT2, BMP6, INHBA and the interaction between TGFβ-signaling and Wnt-signaling by INHBA, INHBE, and FZD8. As expected, genes encoding for collagen synthesis were responsible for ECM organization (Figure 7d). Collectively, these results suggest a gene signature for myofibroblasts following exposure to TGFβ is built upon gene expression patterns in collagen fibril organisation, ECM organisation, cell senescence, cell adhesion, TGFβ signalling, and regulation of pathway restricted Smad phosphorylation.

**Figure 7:**
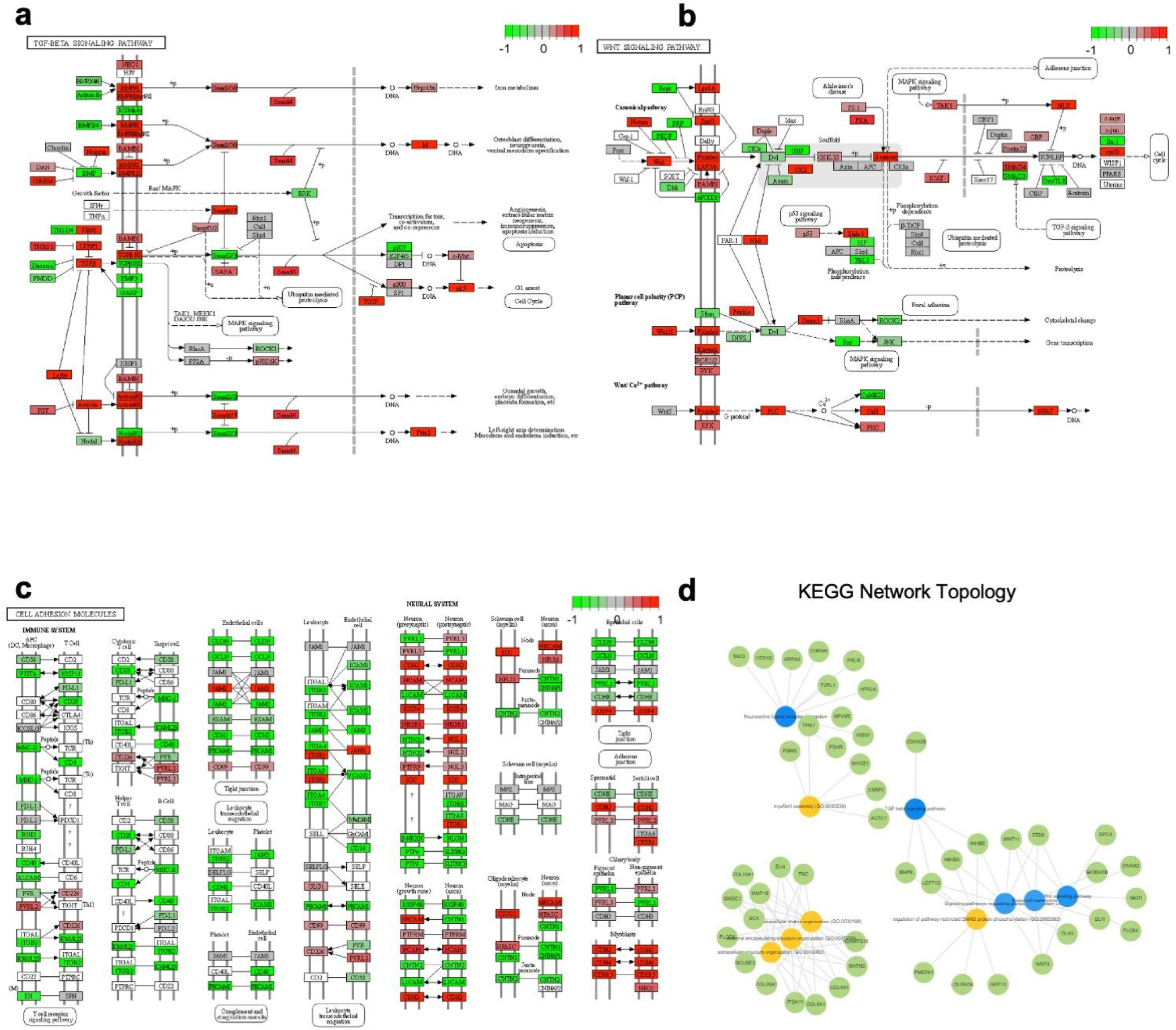
GSEA using KEGG gene set in HTF following TGFβ1 treatment. KEGG pathway map for TGF-B signaling pathway. Rectangles correspond to genes or enzymes plotted by fold changes red = increased fold change, green = decreased, and grey = not available. Colour key gives a range from -1 to 1 and values beyond that range are converted to closest extreme e.g. values >1 converted to 1.**(a)** KEGG pathway map for TGF-B signaling. **(b)** KEGG pathway map for Wnt signaling pathway. **(c)** KEGG pathway map for cellular adhesions molecules pathway. **(d)** Network topology of the top 200 up-regulated DE genes to KEGG pathways.

Next, Gene set enrichment analysis (GSEA) was performed to integrate the DEGs from the TGFβ treatment group in an effort to identify top canonical pathways enriched with publicly available KEGG and GSEA-hallmark gene sets. Myofibroblast differentiation had the highest normalized enrichment score with the highest enriched genes including *ALDH1B1, LOXL2, NEXN, COTL1,* and *C5orf46* (Supplementary table 1)(Figure 8a). Genes encoding for collagen synthesis including *P4HA2* and *P4HA3* were enriched in the collagen fibril organization enrichment plot (Figure 8b) (Supplementary table 1). Tumor suppressors such as *CDKN2B* and oncogenes such as *CCNA1* were highly enriched in cell senescence (Figure 8c) (Supplementary Table 1). *CTGF, TGFβ1, TGFβ3*, and *SMAD7* were highly enriched in the TGFβ-pathway (Figure 8d) (Supplementary table 1). *MYOZ1* was found to be significantly enriched in wound healing organization (Supplementary figure 3a), and *FBLN5* was significantly enriched in extracellular matrix interaction cell adhesion and within the top 20 significant DEGs (Supplementary figure 3b). Families of MMP and collagen genes were highly enriched in extracellular matrix organisation (Supplementary figure 3c). Altogether, our results identified the key genes regulating the myofibroblast transition of HTFs.

**Figure 8:**
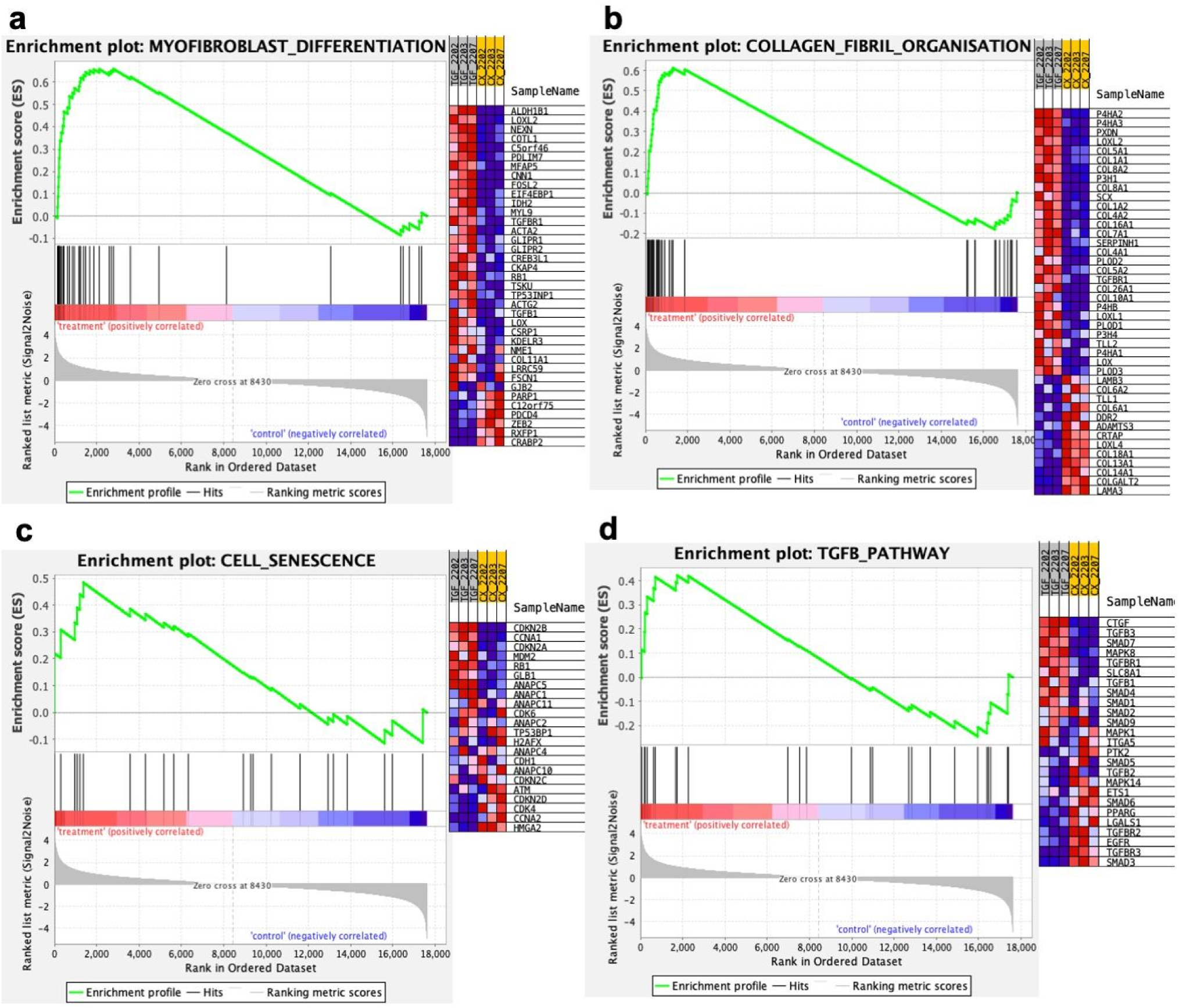
GSEA analysis in HTF following TGF-B1 treatment. Gene set enrichment analysis for TGF-B treatment verse control. **(a)** GSEA plot of myofibroblast differentiation with GSEA standard hallmark gene set with heatmap of gene expression red = upregulated, and blue = downregulated. The enrichment score (ES) is indicated. **(b)** GSEA plot for collagen fibril organisation using GSEA standard hallmark gene set. **(c)** GSEA plot for cell senescence using GSEA standard hallmark gene set. **(d)** GSEA plot for TGFB pathway using GSEA standard hallmark gene set.

## Discussion

This study presents the first RNA-seq study of human Tenon’s fibroblast-to-myofibroblast activation, a critical TGFβ-dependent process involved in postoperative wound healing following glaucoma filtration surgery. Myofibroblast activity drives wound healing both physiologically and pathologically. The progression of fibrotic diseases and cancers is dependent on myofibroblasts that remain pathologically activated rather than undergo apoptosis and vascular pathologies ^9^. TGFβ signalling notably plays a critical role in maintenance of tissue homeostasis, and is hence highly regulated at several steps in both intracellular and extracellular microenvironments ^10^. As this analysis was conducted in myofibroblasts at a 5-day timepoint post-TGFβ treatment, where HTFs expressed a myofibroblastic phenotype, we expected our findings to reflect the direct and indirect effects of TGFβ on activation and inhibitory signalling in myofibroblasts.

Growth factors and reactive oxidants are released from damaged cells and trigger the activation and proliferation of fibroblasts ^11^. CTGF is one such growth factor that has been found to play an important role in TGFβ-dependent myofibroblast differentiation and ECM production, indicated by its upregulated expression being highly enriched in the TGFβ-pathway (Figure 8d). As an important downstream mediator of TGFβ, CTGF has been found to influence myofibroblast differentiation in orbital fibroblasts through increasing of fibronectin and a-SMA protein expression^12^. Hence CTGF may provide a safer target for suppression therapy compared to TGFβ. *ACTA2* appears to be important in myofibroblast contraction and migration. *ACTA2* is involved in the regulation of multiple genes relevant to fibrosis, including *COL1, GFAP, TIMP1, TGFβ*, and *ET1* ^13^. We found *NOX4*, a gene encoding for reactive oxygen species NADPH oxidase 4 enzyme, was upregulated by TGFβ. NOX4-dependent generation of hydrogen peroxide is a requirement for TGFβ induced myofibroblast differentiation, as well as ECM production^14^. *TXNDC5* facilitates proper protein folding in the endoplasmic reticulum and mediates redox reactions via interacting with NADPH oxidase ^11^. These genes promote conjunctival fibrosis by activation of SMAD3-dependent TGFβ-signaling and lead to transdifferentiation of HSCs into myofibroblasts, this results in considerable myofibroblast proliferation and ECM production ^11^. Notably, we observed the downregulated expression of *SOD3* in our TGFβ treated samples. As known ROS scavengers, SODs act to reduce oxidative stress in both extra- and intra-cellular environments, with SOD3 specifically acting in the ECM where it has been found to reduce intracellular ROS levels ^15^. Hence our finding is consistent with findings of SOD3 deficiency contributing to liver fibrogenesis and TGFβ1 mediated EMT ^16^.

TGFβ utilizes components of the ECM such as fibronectin and integrins to communicate and control cell behaviour leading to myofibroblast differentiation. *EDN1, FN1, TNC* and *ITGA11* were upregulated by TGFβ and have been reported in myofibroblast differentiation. *EDN1* has been shown to induce resistance to apoptosis in fibroblasts and contribute to fibrogenesis by the abnormal persistence of the myofibroblast phenotype ^17^. TGFβ can induce expression of fibronectin 1 extra domain A (FN1 EDA) in fibroblasts. *FN1* is not expressed in healthy tissue but is expressed during wound healing, fibroblasts detect FN1 EDA and its presence is required for TGFβ-mediated myofibroblast formation ^18^. *TNC* encodes for tenascin-C a ECM glycoprotein that has been shown to engage integrins to elicit cell specific responses such as fibrotic responses including collagen synthesis and differentiation of myofibroblasts ^19^. *TNC* was upregulated in the TGFβ sample and has been shown to induce myofibroblast differentiation and migration ^20^. Integrins are the main cell-adhesion transmembrane receptors and bind proteins in the ECM such as fibronectin and transduce biochemical and mechanical signals^21^. *ITGA11* was the most significantly upregulated integrin in our data and has been shown to co-localize with a-smooth muscle actin-positive myofibroblasts and was correlatively induced with increasing fibrogenesis in human fibrotic organs. *ITGA11* knockdown has been shown to markedly reduce TGFβ-induced differentiation and fibrotic parameters ^22^. Our findings are consistent with other studies that reported TGFβ upregulation of *ITGA11* and *CTGF* predominantly binding *ITGA11*^23^. We suggest that *ITGA11* is a key regulator of myofibroblast differentiation in HTF and this association may hold potential as a therapeutic target.

Elastic fibre proteins play a pivotal role in guiding and facilitating elastogenesis involved in wound healing. The elastic fibre proteins implicated in this process include fibulins, fibronectins, fibrillins and LTBPs^24^. In our TGFβ sample, we recorded upregulated gene expression of *FBLN5, FBN1, FN1,* and *LTBP1*, alongside downregulated gene expression of *LTBP4* and *FLBN7*. Of particular interest was *FBLN5* which was in the top 20 upregulated DEGs. *FBLN5* encodes the matricellular protein fibulin-5, which performs a vital role in elastogenesis and dysregulated expression of FBLN5 has been reported to occur in pseudoexfoliation glaucoma^25^. These findings correlate with the observed upregulated expression of *FBLN5* being highly enriched in the EMICA pathway.

Typically TGFβ is secreted in a form that is covalently bound to members of the latent TGFβ-binding protein (LTBP) family, observed as a large latent complex. It is via interactions between LTBPs, fibronectin and fibrillin that this complex can be deposited into the ECM ^26^. Hence LTBPs function to regulate the bioavailability of TGFβ, by either localising latent TGFβ in the ECM, or secreting latent TGFβ from cells ^27^. In Lu et al.’s 2017 study of human sclerodermal skin fibroblasts, the knockdown of *LTBP4* resulted in the inhibition of collagen expression through canonical TGFβ/SMAD signalling, while also reducing extracellular levels of TGFβ ^28^. However contradictorily, findings from Su et al.’s recent 2023 *LTBP4* knockdown study in a murine model of renal fibrosis revealed that an *LTBP4* deficiency promoted mitochondrial dysfunction, increased inflammation, oxidative stress via increased ROS production and fibrosis ^29^. Hence we suspect that the regulatory function of LTBP4 is complex and contextual, as it has been found capable of performing both anti- and pro-fibrotic functions. This makes *LTBP4* a prime candidate for further research. In dermal fibroblasts, TGFβ exposure resulted in increased deposition of fibrillin-1 and fibronectin into the ECM as a consequence of suspected myofibroblast activation ^30^. This associates our observed upregulation of *FBN1* and *FN1* in our TGFβ treated HTFs with myofibroblast activation.

TGFβ mediated epithelial to mesenchymal transition is important for myofibroblast transition. Epithelial to mesenchymal transition causes several changes to cellular phenotype and cytoskeleton remodelling and increased cell migration. We identified several important genes interacting with TGFβ for this process. *NREP* was upregulated in the TGFβ sample, and regulates the expression of TGFβ1 and has been shown to regulate myofibroblast differentiation. *NREP* is thought to stimulate the expression of TGFβ1 by the methylation of NREP promoter and activating TGFβ1 5’/3’ UTR ^31^ . Knockdown studies of *SCUBE3*, which was upregulated in the TGFβ sample, have implicated its involvement with lung cancer tumorigenesis and cancer metastasis. SCUBE3 binds to TGFβ type 2 receptor and activates TGFβ signalling triggering epithelial-mesenchymal transition ^32^. *DACT1* was upregulated in the TGFβ sample; it has been shown to inhibit the Wnt signalling pathway. Therefore, DACT1 may release inflammatory factors that were previously suppressed by Wnt and promote local inflammation ^33^. Of the Wnt signalling pathways that were highly enriched in our gene set, the Wnt/Ca^2+^ pathway (Figure 7b) is known to play a critical role in profibrotic and proinflammatory processes including actin polymerisation, cell adhesion, cell migration and NFAT signalling ^34^. A key component of this pathway involves the activation of the serine-threonine phosphatase enzyme calcineurin (CaN). RCAN2, encoded by *RCAN2*, has been identified as an inhibitor of the Wnt/Ca^2+^ pathway by binding CaN and subsequently inhibiting CaN’s protein phosphatase activity^35^. Findings from our GO analysis are in line with these previous findings as the molecular function of *RCAN2* was found to be involved in protein phosphatase regulator activity (Figure 6e, GO:0019888). Hence we suspect that our observation of *RCAN2*’s downregulated expression in the TGFβ sample is associated with myofibroblast activation through promotion of the Wnt/Ca^2+^ signalling pathway.

The regulation of cell cycle process and cell senescence pathways were significantly enriched and largely driven by genes *CDKN2, CCNA1, CDKN2A, CDKN2D,* and *CDK4*. TGFβ causes G1 phase cell cycle arrest and CDKN2B complexes with CDK4 to prevent CDK4 activation and producing cell cycle arrest ^36^. These genes are known cell cycle regulators and CDKN2b/p15 has been reported to take part in multiple pathologies such as primary open angle glaucoma and cardiac fibrosis^36^. Interestingly, TGFβ treatment resulted in downregulated expression in the majority of cell cycle and proliferation marker genes (Figure 5). The only exception to this observation was the upregulated expression of both the tumour suppressor gene *CDKN2B* (Figures 4b and 4c) and the cell cycle proliferator gene *TMPO* (Figure 5). Lung carcinogenesis has been linked to upregulated expression of *TMPO* ^37^, while knockdown of *TMPO* in glioblastomas has been found to inhibit apoptosis-dependent cell proliferation and arrest cell cycle progression at the G2/M Phase ^38^. Collectively these findings are consistent with other studies reporting the role of TGFβ as a tumour suppressor and inhibitor of cell proliferation, such that cell cycle arrest and apoptosis evasion occur ^39^.

*CDK6* is a known cell cycle regulator for G1 to S phase transition. Dysregulated expression of *CDK6* has been reported in a variety of malignancies. Palbociclib is a commercially-available selective inhibitor of *CDK6* which has shown significant inhibition of myeloproliferative neoplastic progenitors and cells, and amelioration of bone marrow fibrosis in myelofibrosis ^40,41^. These findings are consistent with other studies and highlight the biological effects of TGFβ at the cellular level.

*TGFβ1* and *TGFβ3* were upregulated in the TGFβ treatment group. The 3 TGFβ isoforms have highly homologous receptor-binding domains and similar effects on target cells but with divergent primary amino acid sequences in the latency-associated peptide (LAP) domains. Isoform-specific function may be related to different expression in cell types and different extracellular environments ^42^. *TGFβ1* was enriched in pathways such as TGFβ-pathway, myofibroblast differentiation, and collagen fibril organisation whereas *TGFβ2* and *TGFβ3* were not. This suggests that HTF induced fibrosis is largely a *TGFβ1* driven process. *TGFβ3* appears to play a role in regulation of epithelial to mesenchymal transition and TGFβ-pathway rather than myofibroblast differentiation which is consistent with previous reports of TGFβ3 expression being restricted to mesenchymal cells^42^.

Our findings highlight the important molecular mechanisms of TGFβ-signaling, in particular positive and negative regulation of pathway-restricted Smad protein phosphorylation (Figure 6c; GO:0060393 & GO:0060394). The main genes upregulated in this biological process were *GDF10, PMEPA1, LDLRAD4, INHBA, LEFTY2, BMP6* and *SMAD7*. Canonically TGFβ signal transduction occurs via activation of downstream mediators Smad2 and Smad3, which are negatively regulated by Smad7 as part of a negative feedback loop ^43^. Smad7 inhibits phosphorylation of R-Smads(2/3) by competitively binding TβR-I activated by TβR-II ^44^ . We speculate that the upregulated expression of *SMAD7* is associated with the upregulated expression and enrichment of the *SMURF1* gene as their encoded proteins regulate each other’s functions. In order for Smad7 to perform its inhibitory function it is required to interact with Smurf1 to be exported from the nucleus to the cytoplasm ^45^. *INHBA* encodes inhibin beta-A, which functions as a subunit of activin A and is classified as a ligand of the TGFβ superfamily^46^. INHBA attenuation in renal fibroblasts has been found to significantly reduce their expression of profibrotic markers, inhibiting cell migration and proliferation ^47^. To promote the TGFβ signalling pathway, INHBA is able to homodimerise and bind the activin type I/II receptor complex, which then can phosphorylate R-Smad proteins ^48,49^. Hence, we suspect that upregulated INHBA expression correlates with upregulated expression of *ACVR1* (encodes type I receptor, ALK2) and *ACVR2B* (encodes type II receptor) to promote Smad-dependent TGFβ signalling.

*PMEPA1, LEFTY2,* and *LDLRAD4* are all negative regulators of TGFβ-signaling. *PMEPA1* and *LDLRAD4* are part of the PMEPAI-family genes and encode transmembrane proteins that have been found to facilitate TGFβ signalling inhibition by competitively targeting SARA through sequestration of the Smad2/3 protein complex and attenuating R-Smad complex recruitment to the TβR-1 for phosphorylation ^50,51^. Once activated by TGFβ, *LEFTY2* inhibits the phosphorylation of Smad2 and downstream heterodimerisation Smad4. Lefty also regulates the extracellular matrix components and inhibits the profibrotic effects of CTGF ^52–54^.

While the results achieved in this study advanced our understanding of the molecular mechanisms of ocular fibrosis following GFS, this study has its limitations. Although current literature supports RNA-seq analysis of HTF as a homogenous population, single cell RNA-seq would enable high resolution transcriptome analysis and potentially identify heterogeneity or subtypes within the HTF and myofibroblast population. This high resolution transcriptome analysis could then lend to the identification of tissue biomarkers for ocular fibrosis, and together these tools could be instrumental in performing detailed clinical phenotyping. This would enable clinicians to not only identify groups of patients that are likely to scar more severely than others, but also work toward developing a tiered approach to antifibrotic therapy that is more personalised to the patient. Another limitation of the study was that specimen collection was performed at the time of GFS, from patients who were receiving medical therapy for glaucoma (Supplementary Table 2). In using these glaucoma medications, most, if not all the patients were exposed to the preservative benzalkonium chloride, which is frequently associated with adverse reactions ^55^. Recorded adverse side effects of benzalkonium chloride include but are not limited to trabecular meshwork cell apoptosis, conjunctival and anterior chamber inflammation, corneal cytotoxicity, tear film instability, cataract development and macular oedema ^56–58^. Yet the effects of these medications on gene expression profile remain largely unknown. The small sample size of this study was also a limitation. Although a sample size of 3 patients is within current practice for genomic studies a larger sample size with high quality reads would increase the statistical power of the findings. Future knock out or knock down studies would be useful to validate the DEGs identified in this study, such that their suitability as targets for drug or gene therapy can be assessed.

In investigating a model of HTF transition into myofibroblasts at the transcriptomic level, we gained greater insight into the novel molecular interactions and causal pathways of ocular fibrosis. This study establishes a gene signature of myofibroblast activation from HTFs that is uniquely characterised by genes involved in regulation of myofibroblast differentiation, collagen fibril organization, cell cycle arrest, TGFβ-signaling pathways, and wound healing organization. The results of this study establish an essential milestone for the future development of effective antifibrotic therapies targeting scar tissue formation following GFS.

## Methods

### HTF Isolation

Isolation of HTFs was conducted in compliance with guidelines approved by the institutional ethics committee (Eye and Ear Hospital Human Research Ethics Committee project no. 16/1294H). Following explanation of the nature of the study, informed consent was obtained from patients. HTFs were propagated from explanted subconjunctival Tenon’s capsules collected during GFS performed in three patients and handled in accordance with the tenets of the Declaration of Helsinki.

### Cell Culture

HTFs were maintained in Dulbecco’s Modified Eagle Medium (DMEM; Sigma-Aldrich) containing Gibco Penicillin–Streptomycin– Glutamine (Thermo Fisher Scientific) and 10% fetal calf serum (FCS; Cytiva, Marlborough, MA) at 37°C with 5% CO_2_ in 50 mL and 160 mL cell culture flasks (Greiner Bio-One). Human Tenon’s Fibroblasts were passáged when flasks achieved 80% confluency by dissociating monolayers with a Trypsin-EDTA solution (0.25%; Sigma-Aldrich). Fibroblasts obtained on passáges 6 through 12, after initial cultures were established, were used. Phenotype verification of the cultured HTFs was achieved with the use of anti–fibroblast-specific protein 1 (S100A4) antibody (1:100; ABF32; Merck Millipore, Bayswater, VIC, Australia)(Supplementary Figure 1). HTFs (approx. 40,000 cells/well) were seeded onto coverslips in 6-well Nunc cell culture plates (Thermo Fisher Scientific). Then HTFs were serum-starved in DMEM with 1% FCS overnight in preparation for treatment. HTFs were then assigned to either the control (medium alone) or the TGFβ1 (10 ng/mL) treatment group. Following treatment, HTFs were fixed with 4% paraformaldehyde (PFA; P6148; Sigma-Aldrich, CA, USA) over a range of time-points (from Days 3-14).

### Immunocytochemical/ Immunofluorescence Staining

Following fixation HTFs were rinsed with Dulbecco’s Phosphate Buffered Saline (DPBS; Gibco) and permeabilised with 0.1% Triton X-100 (Sigma-Aldrich, St Louis, MO, USA) for 5-10 minutes and rinsed again with PBS. In order to prevent non-specific binding of the primary antibody, a blocking solution, UltraVision Protein Block (TA-125-PBQ; ThermoFisher Scientific, CA, USA), was applied to coverslips for 30 mins. HTFs on coverslips were then left to incubate overnight at 4°C with the primary anti-’α-SMA antibody (M0851; Dako, CA, USA). Following incubation period, HTFs on coverslips were rinsed thoroughly with PBS and were treated with a fluorophore-labelled secondary antibody, Alexa Fluor 568 goat anti-mouse IgG (H+L) (A11004; Invitrogen, OR, USA), for 1 hour in the dark at room temperature (22°C), then rinsed again with PBS. Nuclear counterstaining was conducted with the application of 4′,6-diamidino-2-phenylindole (DAPI) (1:1000; ab228549; Abcam, Cambridge, UK) for 10 mins in the dark at room temperature. Finally coverslips were mounted onto microscope slides with DPX mounting media (20242; Labworks, Knox City Centre, VIC, Australia). All slides were observed at 20x magnification using a fluorescence microscope (Axio Scope.A1 with Axiocam 105 colour; Zeiss, Oberkochen, Germany) and analysed with corresponding software (Zen 3.1- blue edition, Zeiss, Oberkochen, Germany).

### Real Time qPCR

RNA extraction was performed using illustra RNAspin Mini Kit (GE Healthcare Life Sciences), following manufacturer’s instructions. RNA concentration and quality were measured using SimpliNano (GE Healthcare Life Sciences). cDNA was synthesised by reverse transcription of RNA. RT-qPCR was performed using TaqMan^TM^ Fast Advanced Master Mix (4444554; Thermo Fisher Scientific, Lithuania) and Taqman gene expression assay probe for *ACTA2* (Hs00426835_g1) and the housekeeping gene 18s rRNA control mix (4318839; Applied Biosystems, Kingsland Grange, Woolston, Warrington, UK). RT-qPCR was performed on QuantStudio™ 7 Flex Real-Time PCR System (Thermo Fisher Scientific, Scoresby, VIC, Australia), following manufacturer’s instructions. The delta delta Ct (ΔΔCT) method was used to calculate and compare relative mRNA levels to control. Data were expressed as mean ± standard deviation (SD) and were analyzed with two-way analysis of the variance (ANOVA) followed by post hoc Šídák’s multiple comparisons test. A value of P < 0.05 was regarded as statistically significant.

### RNA Sequencing

RNA was extracted using the Illustra RNAspin Mini Kit (GE Healthcare Life Sciences) according to manufacturer’s instructions. RNA quality was checked by bioanalyzer, followed by library construction using the TruSeq Stranded mRNA kit (Illumina) and sequenced using Novaseq 6000 (Illumina) with 100bp single-end sequencing, at a depth of ∼20 million reads per sample (Australian Genome Research Facility).

### Bioinformatic analysis

Following the abundance estimates of transcripts generated by *Salmon* v1.8, the pseudocounts were mapped to the GRCh38 genome assembly using the *tximport* v1.22.0 package^59^. The gene count matrix was inputted as an DESeq2Dataset object using the *DESeqDataSetFromTximport* function, then the DESeq2Dataset object was normalised using the *counts* function to make fair gene expression comparisons between samples^60^. The normalised dataset was analysed with the *DESeq2* v1.34.0 R package using rlog transformation. The sample-level quality assurance and distribution bias was measured using principal component analysis while the gene-level QC was performed using hierarchical clustering. For differential expression analysis, the significant differentially expressed genes were determined using the *filter* function with adjusted p value of < 0.05 and fold change > 2.0. The expression data of significant differentially expressed genes was visualised using the *ggplot2* v3.3.6, *pheatmap* v1.0.12 and *EnhancedVolcano* v1.12.0 R package^61–63^. GO enrichment analysis was performed to investigate relationships between significantly expressed genes and their cellular component, biological process, and molecular function. Only terms with a p-value <0.01 were considered significant. Network topology analysis were performed using Enrichr with the top 50 upregulated DEGs ^64–66^. Gene set enrichment analysis was performed using GSEA 4.3.2 and MSigDB 2023.1 (UC San Diego and Broad Institute) ^67,68^ a false-discovery rate of q-value <0.25 and a p-value <0.01 were considered significant. The MSigDB gene sets genes known to be significant in the following pathways: cell cycle phase and proliferation, TGFB signaling, Wnt signaling, NFKb pathway, fibroblast markers, ECM organization, wound healing regulation, myofibroblast differentiation, collagen fibril organization, cell cycle regulation, and ocular fibroblast markers. Further GSEA and pathway analysis was conducted using msigdbr ^69^ and clusterProfiler packages ^70^. KEGG network topology for the top 50 DEGs was performed using Enrich to obtain gene and KEGG term interactions ^65^.

## Data availability

The transcriptome data generated in this study are available in the NCBI Gene Expression Omnibus database (GSE accession pending), including raw data and processed data.

## Author contribution

Conceptual design: ECC, RCBW, JG; Conduct experiments: ZP, RK, JG; Data analysis: AB, RCBW, ECC, JG, ZP; Funding: JG, RCBW; Manuscript writing & review: ZP, AB, RCBW, ECC, JG. All authors approved the manuscript.

## Acknowledgement

This work was supported by the Royal Victorian Eye and Ear Hospital Early Research Career Support Grant (JFG). ZP and AB are supported by an Australian Government Research Training Program Scholarship. RCBW is supported by the University of Melbourne, and the Centre for Eye Research Australia, National Health and Medical Research Council (GCT1184076) and Medical Future Research Fund (MRF2024365). The Centre for Eye Research Australia acknowledges the Victorian State Government’s Department of Innovation, Industry and Regional Development’s Operational Infrastructure Support Program.

## Supplementary figures and tables

**Supplementary table 1:**
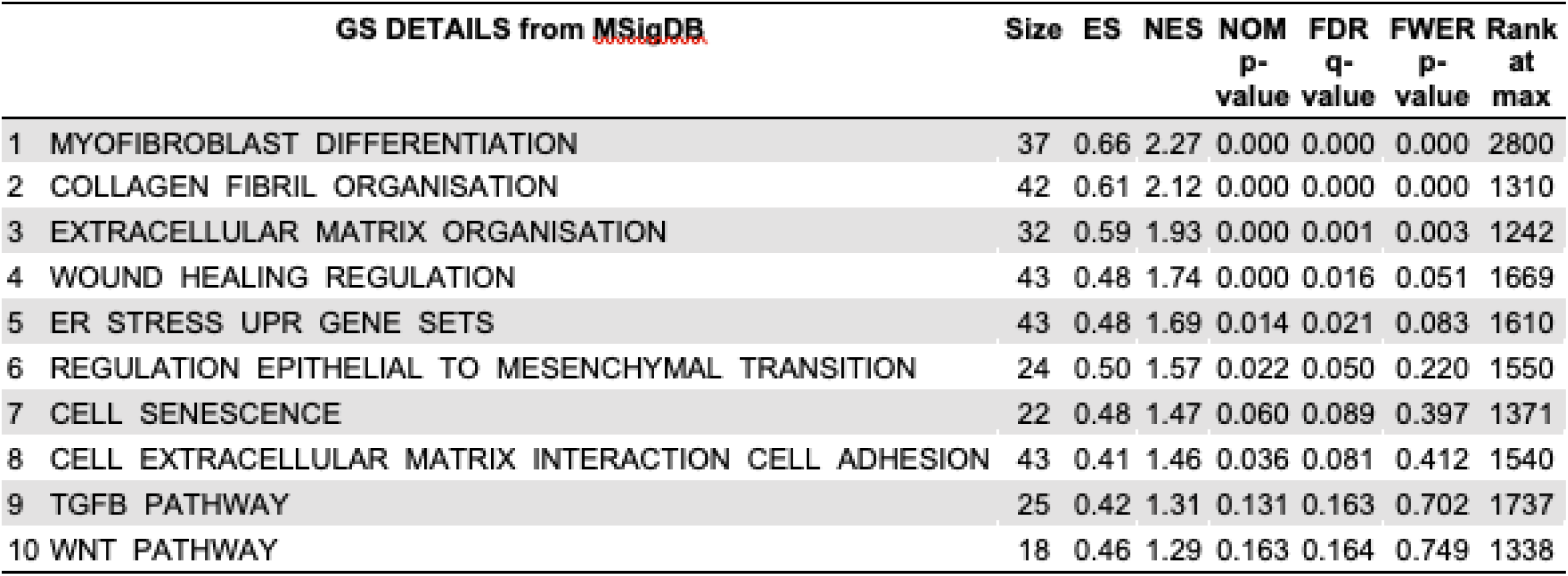
GSEA table for TGF-B1 treatment compared to control. GSEA enrichment results for the top 10 enrichment terms. ES: enrichment score, NES: normalized enrichment score, NOM p-value: nominal p-value, FDR q-value: false-discovery rate q-value, FWER p-value: familywise-error rate p-value.

**Supplementary table 2:** Prescribed glaucoma medications used by patients. Glaucoma medications being used by patients at the time of tissue biopsy collection.

| **Glaucoma Medications** | **Method of Application** | **Preservative** | **Mechanisms of action** |
| --- | --- | --- | --- |
| Bimatoprost | Topical | Benzalkonium chloride | Prostaglandin analog |
| Azarga<br>(brinzolamide/timolol fixed combination) | Topical | Benzalkonium chloride | Carbonic anhydrase inhibitor<br>β-blocker |
| Brimonidine | Topical | Sodium chlorite | α2-adrenergic agonist |
| Simbrizina<br>(brinzolamide/brimonidine fixed combination) | Topical | Benzalkonium chloride | Carbonic anhydrase inhibitor<br>α2-adrenergic agonist |
| Xalacom<br>(timolol/latanoprost fixed combination) | Topical | Benzalkonium chloride | β-blocker<br>Prostaglandin analog |
| Prednifrin forte | Topical | Benzalkonium chloride | Glucocorticoid |
| Fluoromethalone | Topical | Benzalkonium chloride | Corticosteroid |
| Diamox (acetazolamide) | Oral | N/A | Internal carbonic anhydrase inhibitor |

**Supplementary figure 1:**
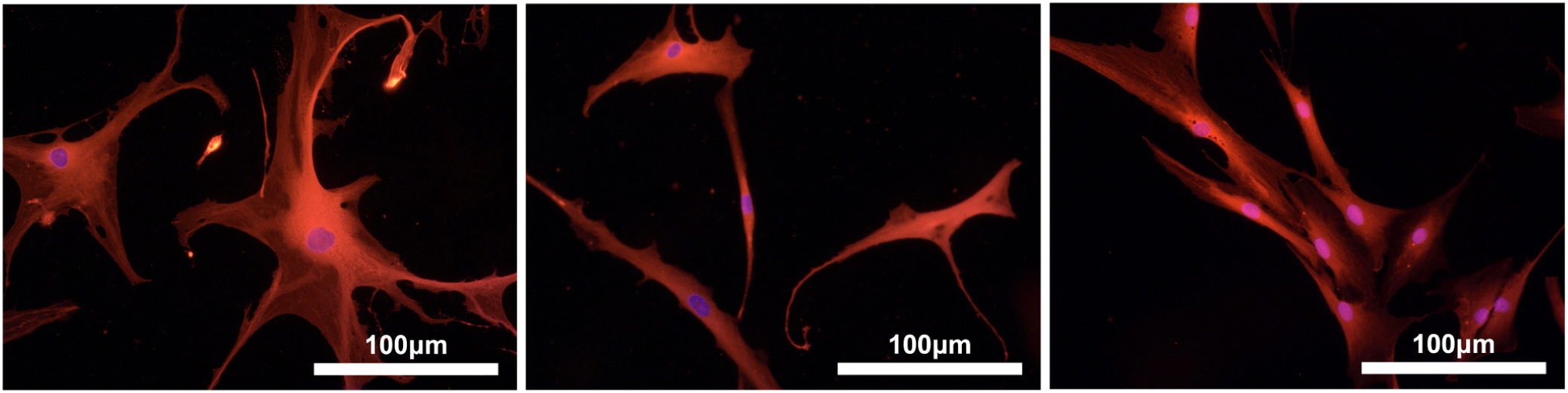
Characterisation of Human Tenon’s fibroblasts (HTFs) Cultured primary HTFs were stained with S100A4 (red) and nuclei counterstained with DAPI (blue). Fibroblast phenotype verified by cells presenting red-stained cytoplasm. (N=3)

**Supplementary figure 2:**
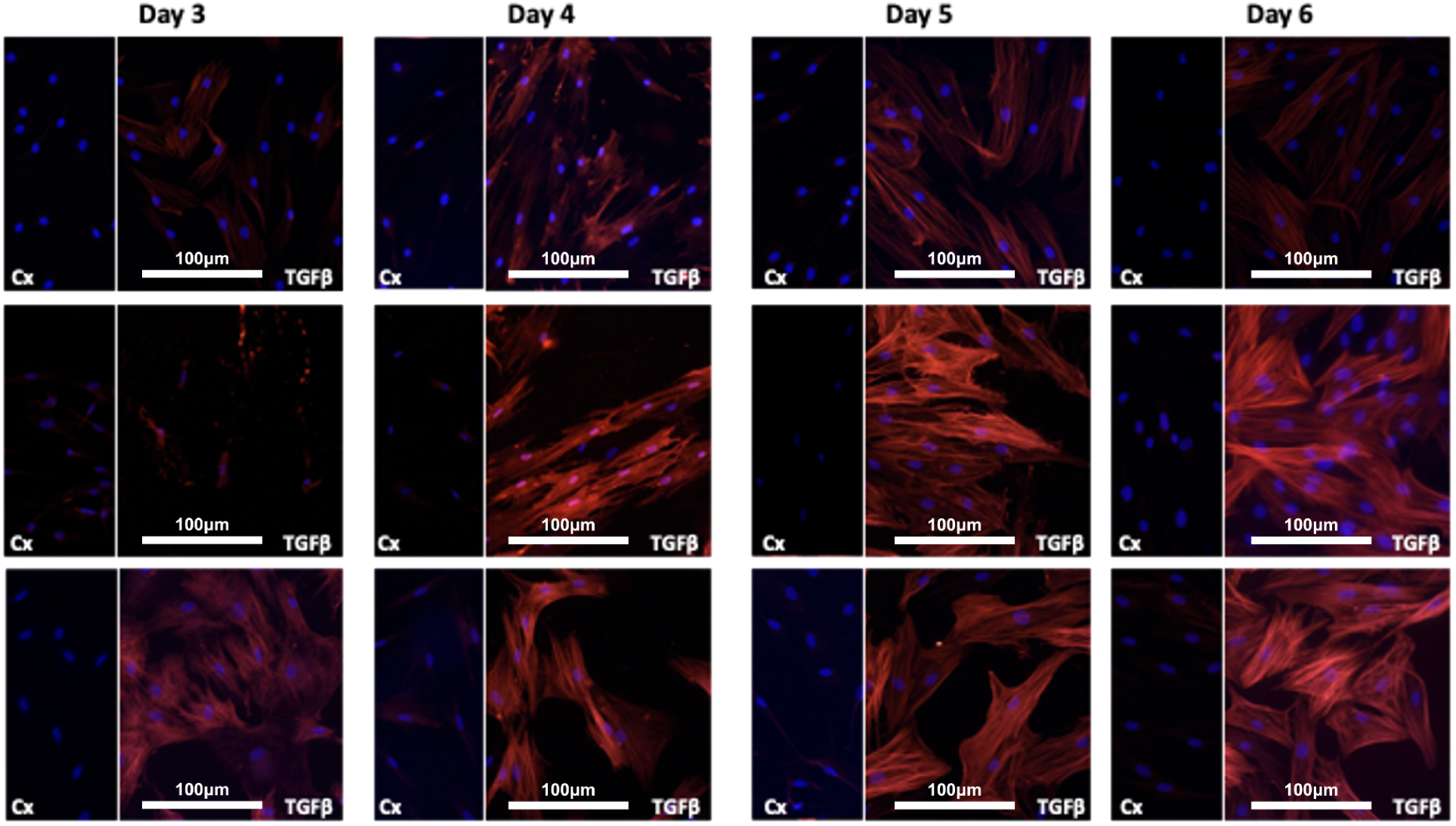
Time-Point Study Triplicate Phenotypic Imaging of HTFs after 3-5 days of TGFβ Treatment. Cultured primary HTFs were stained with αSMA-antibody and nuclei counterstained with DAPI (blue). Red-stained fibres indicate de novo expression of αSMA in collagen stress fibres of myofibroblasts. Cx= Control sample; TGFβ = Transforming Growth Factor Beta treated sample (N=3)

**Supplementary figure 3:**
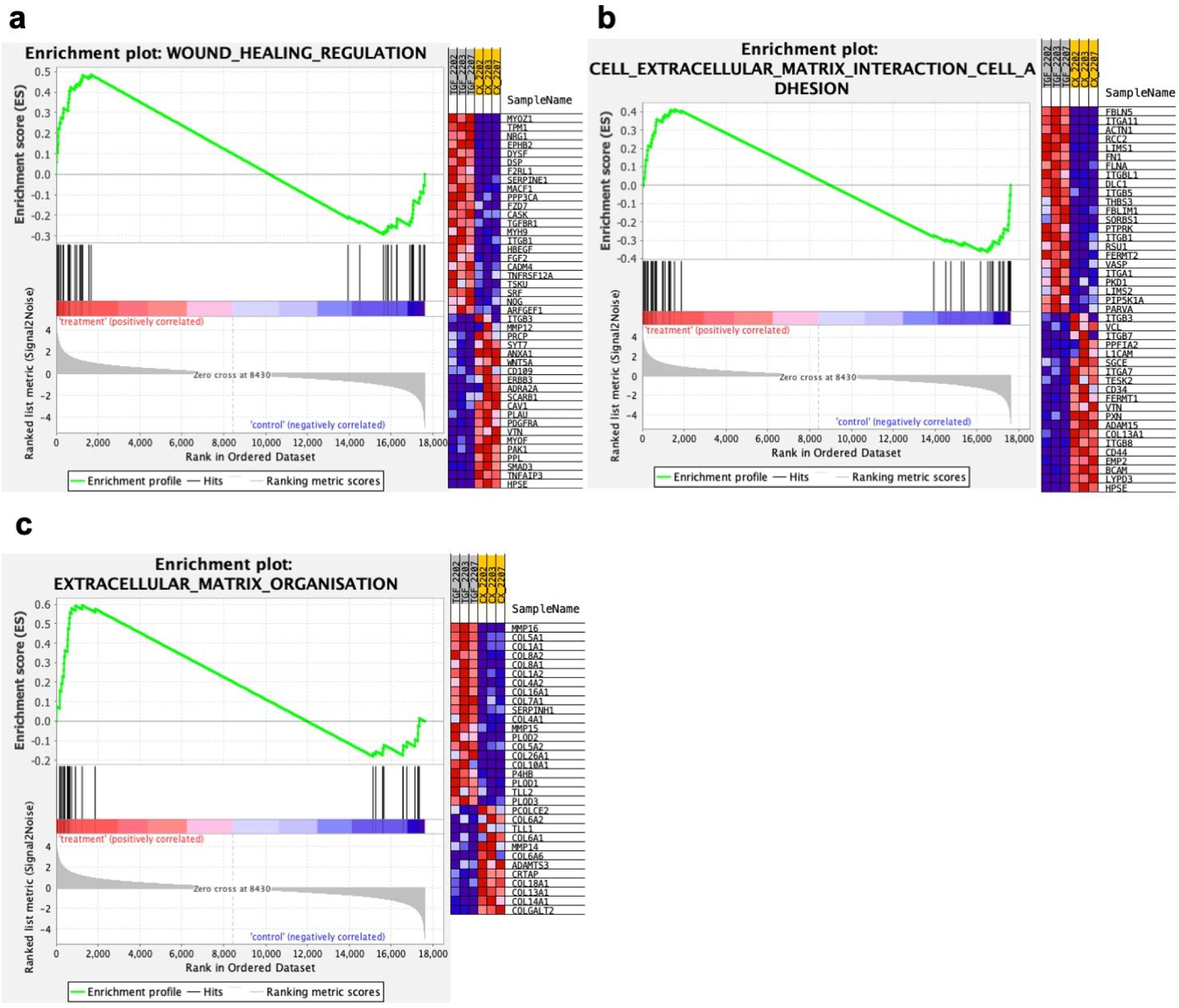
GSEAanalysis for TGF-B treatment versus control. **(a)** GSEA plot of wound healing regulation with GSEA standard hallmark gene set with heatmap of gene expression red = upregulated, and blue = downregulated. The enrichment score (ES) is indicated. **(b)** GSEA plot for cell extracellular matrix interaction cell adhesion using GSEA standard hallmark gene set. **(c)** GSEA plot for extracellular matrix organization using GSEA standard hallmark gene set.

