## Supplementary figures and images for "Transcriptomic analysis of TGFβ-mediated fibrosis in primary human Tenon’s fibroblasts"

### Supplementary Figures and Tables

#### Slide 1
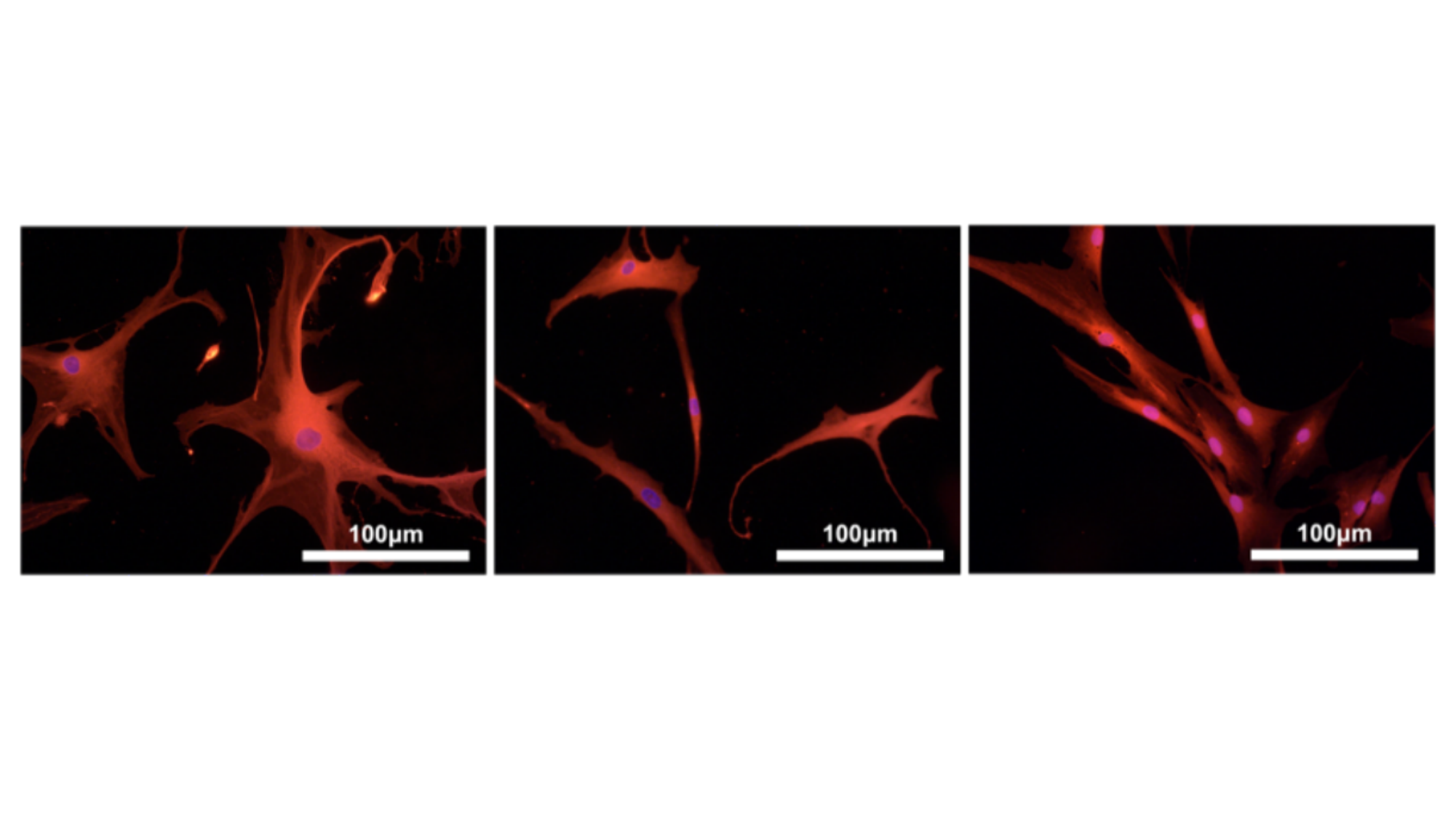

#### Slide 2
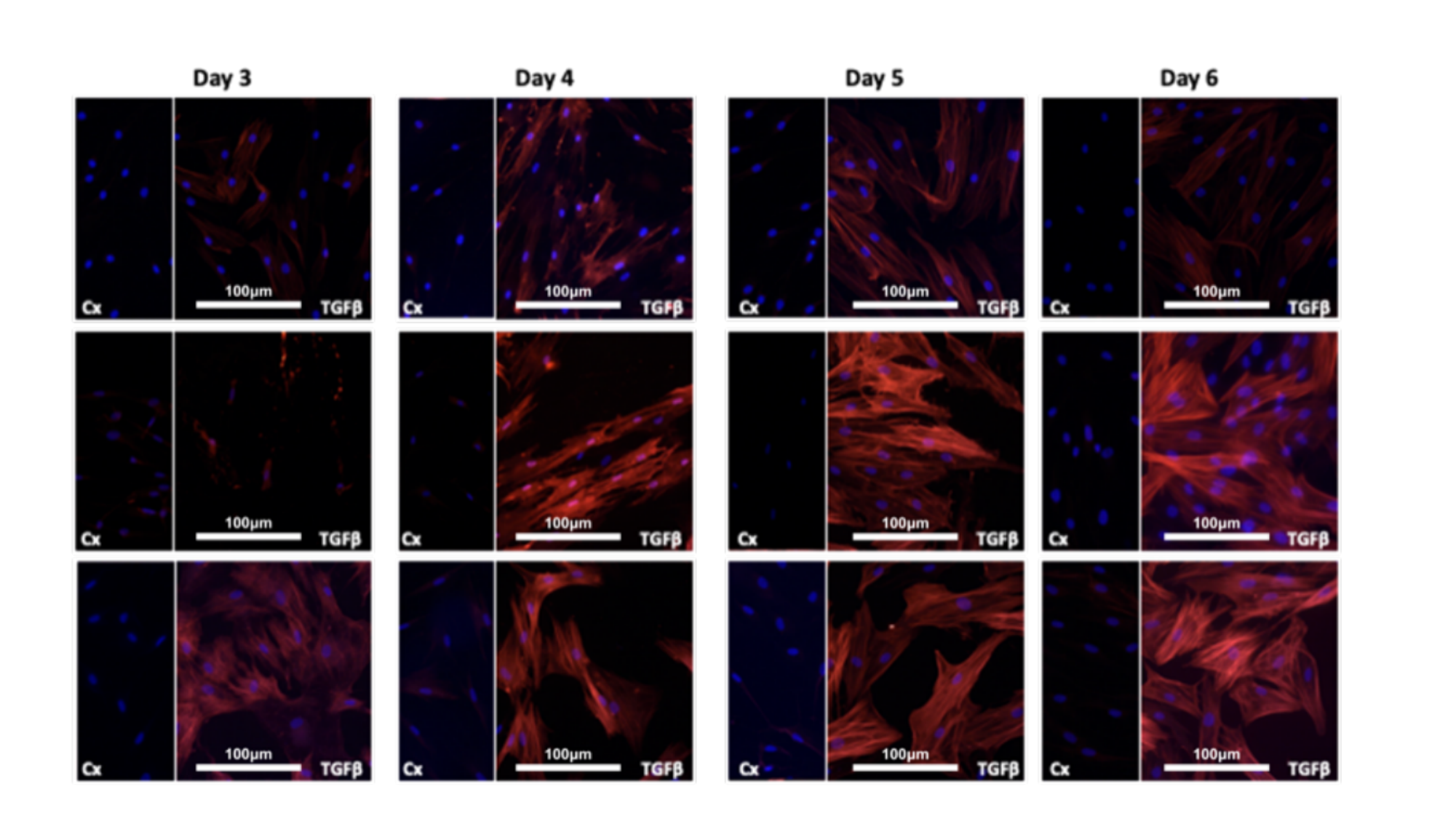

#### Slide 3
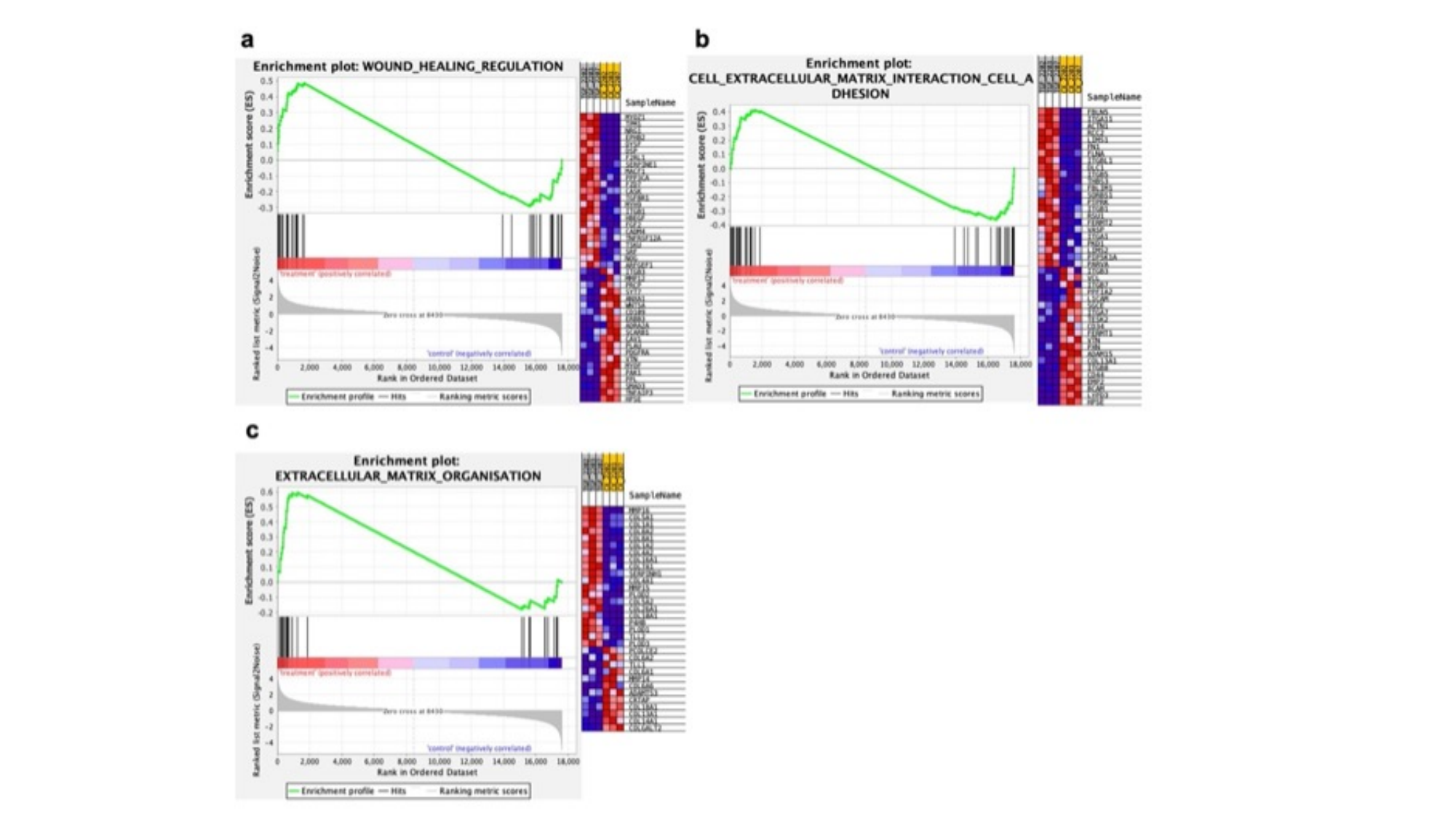

#### Slide 4
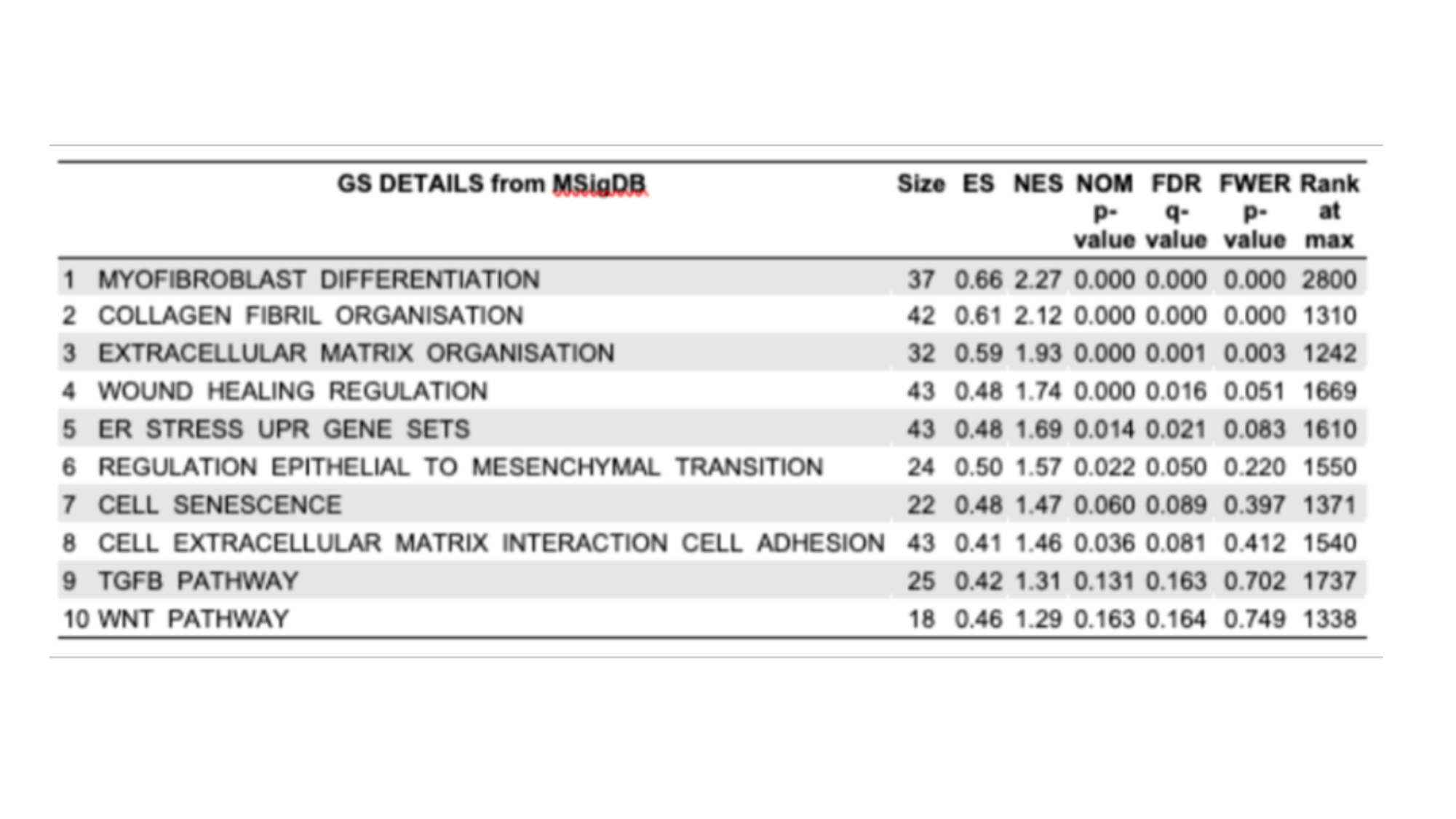

#### Slide 5
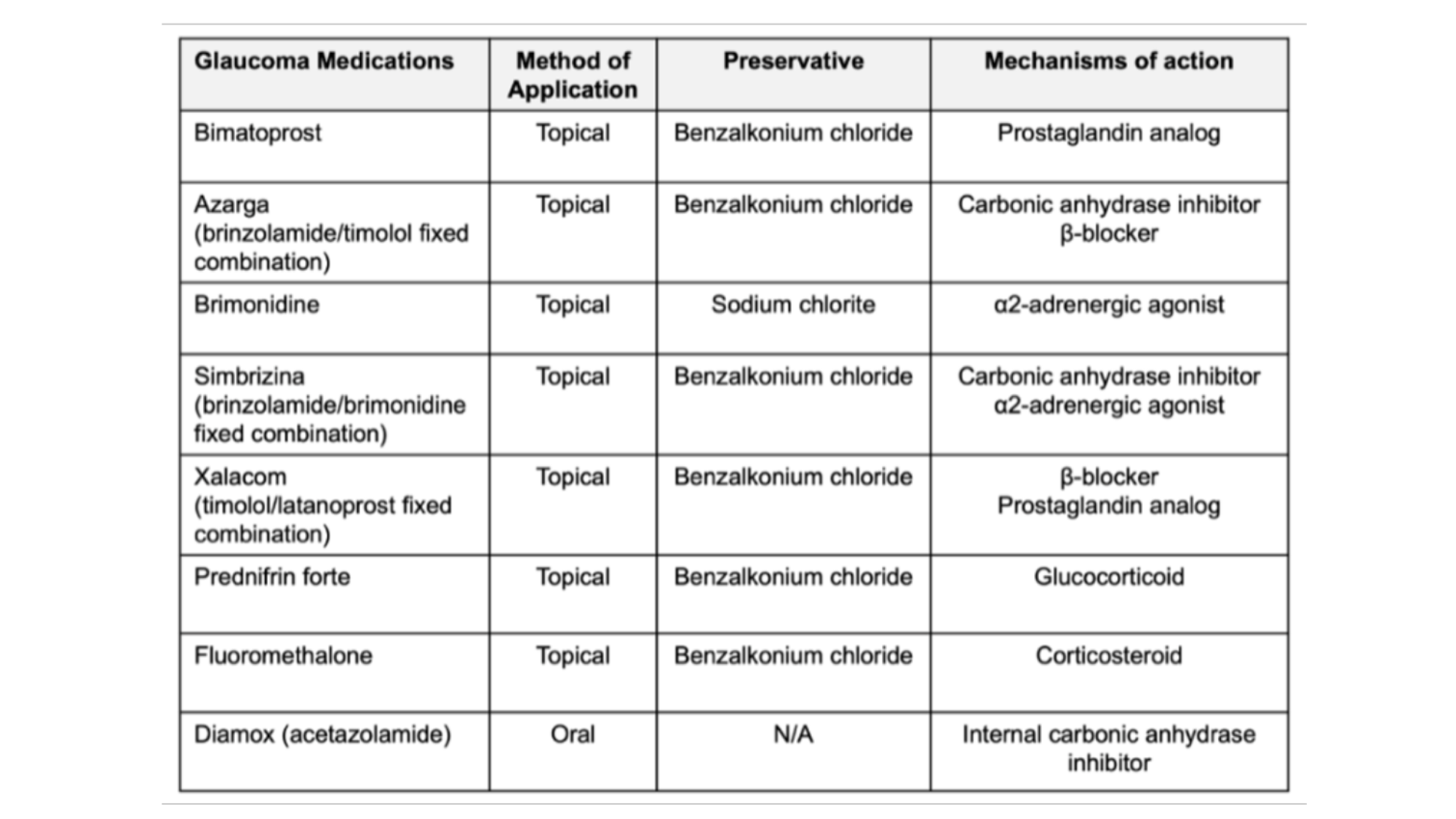
